# Western diet reversibly alters the olfactory mucosa and impairs the response to appetitive food cues

**DOI:** 10.1101/2025.05.16.654424

**Authors:** Louise Eygret, Enisa Aruci, Vincent Canova, Julie Paradis, Vanessa Soubeyre, Tibor Chomel, Rémi Capoduro, Xavier Grosmaitre, Xavier Fioramonti, David Jarriault

## Abstract

Current feeding behaviors contribute to the epidemic levels of obesity and diabetes observed in Europe and worldwide. Together with other sensory modalities, olfaction is involved in the control of food intake. Olfactory cues can influence eating behaviors, yet the nutritional status and diet can also alter olfactory abilities. Patients with metabolic disorders present impaired olfactory sensitivity which could in turn worsen their eating behaviors. Here we examined the short-term impact of a Western diet enriched in fat and sugar (High-Fat High-Sugar, HFHS) on the anatomy and physiology of the olfactory epithelium of mice. After 8 weeks of diet, HFHS fed animals presented higher adiposity without overweight, were glucose intolerant without any change in basal blood glucose or plasma insulin. A buried food test indicated impaired olfactory capacities in the HFHS group. Whereas food related odours increased food intake in control chow fed animals, HFHS mice showed an altered response to olfactory appetitive food cues. HFHS fed mice presented olfactory sensory neurons (OSN) with shorter cilia. Finally, electro-olfactogram (EOG) recorded in response to different odorant molecules showed lower amplitudes in HFHS fed mice. HFHS diet withdrawal during one month at the end of the HFHS diet exposure improved metabolic parameters and restored both the OSN cilia length and EOGs. Our results show that diet enriched in fat and sugar can rapidly alter the physiology of the olfactory epithelium before the development of significant metabolic disorders. Anatomical changes of individual olfactory sensory neurons may participate to the reduced olfactory sensitivity.

## Introduction

Olfaction is a critical sensory modality contributing to essential aspects of life including feeding behaviour, hazard avoidance and social interactions. Olfaction and nutrition are deeply interrelated. The olfactory sense is involved in the detection of food, its hedonic appreciation and the regulation of appetite. Conversely, nutritional state affects olfactory sensitivity (Cameron et al., 2012; O’Doherty et al., 2000; Stafford & Welbeck, 2011), with satiety inducing decreased sensitivity of the olfactory sense in contrast to a higher sensitivity observed after fasting. Decreased olfactory abilities have repeatedly been associated with diet induced obesity in humans (Simchen et al., 2006; Skrandies & Zschieschang, 2015). Higher body mass index correlated with poorer olfactory sensitivity. Children with simple obesity demonstrate significantly lowered thresholds of detection and of identifying odours (Obrębowski et al., 2000). Type 2 diabetes is often associated with olfactory dysfunction (Le Floch et al., 1993; Sienkiewicz-Oleszkiewicz & Hummel, 2024). Similarly, long-term western diet exposure-induced obesity has been associated with impaired olfactory abilities in animal models (Chelette et al., 2022; Lietzau et al., 2020; Takase et al., 2016; Thiebaud et al., 2014; Zou et al., 2022).

Mechanisms have been explored in rodents showing the sensitivity of the olfactory system to metabolic hormones such as leptin, insulin or ghrelin (reviewed in Palouzier-paulignan et al., 2012). Numerous receptors of metabolic peptides or hormones are expressed in the olfactory mucosa and in the olfactory bulb, making them opportune sites for a metabolic modulation at short- and long-term. Perfusions of the olfactory mucosa with the two anorectic factors insulin or leptin decreased the electroolfactogram (EOG) response in mice (Savigner et al., 2009) whereas treatment of nasal mucosa with the two orexigenic factors ghrelin and adiponectin increased the number of activated olfactory sensory neurons (OSNs) (Loch et al., 2013, 2015). Diet composition namely the nature of the lipids consumed was shown to impact the physiology of the OSNs. Western diets composition has changed over the last decades with higher amounts of saturated fat and n-6 polyunsaturated fatty acids (PUFAs) and small amounts of n-3 PUFAs (Blasbalg et al., 2011; Cordain et al., 2005; Sheppard & Cheatham, 2018). Deficiency in n-3 PUFAs was shown to reduce the EOG response of the olfactory mucosa in association with an impairment of the expression of genes involved in olfactory signalling (Soubeyre et al., 2023a).

Here we tested the effect on the peripheral olfactory system of a short-term high fat high sugar diet exposure for 8 weeks which does not induce increased body weight and the capacity of food related odorant to modulate food intake. We also tested whether HFHS reversal to standard chow for 4 weeks reversed impairments of the olfactory system. Thus, 8-weeks HFHS diet induced a change in the structure of olfactory sensory neurons which was associated with impaired olfactory abilities. Diet reversal to standard chow restored the olfactory cilia length and the olfactory capacities. Finally, our data show that HFHS diet abolished the appetitive response to food odours highlighting the drastic impact of olfactory alterations on feeding behaviour.

## Materials and methods

### Animals

All procedures described in this study were approved by the local ethics committee and conducted in accordance with the European guidelines for the care and use of laboratory animals (2010/63/EU) and were approved by the French Ministry of Higher Education, Research and Innovation, the local ethical committee of the University of Bourgogne (APAFIS #01286) and the University of Bordeaux (APAFIS#34196). Transgenic male mice were bred and housed 2-3 per cage at our local animal care facility with free access to water and food. All experiments were conducted on 5 to 16-week-old C57BL/6J male mice reared in an inverted 12L:12D lighting schedule (lights on at 5pm). Animals were fed for 8 weeks either a standard diet (A03, SAFE, France) or a diet enriched in fat and sugar (HFHS diet; 46% energy from fat and 38% energy from carbohydrates; 235HF, SAFE, France). In the reversal protocol, a group of mice were fed with the standard diet for 4 weeks after the 8 weeks of HFHS diet.

### Glucose tolerance, Indirect calorimetry, Fat mass evaluation, and locomotor activity

Glucose tolerance test: Animals were housed in individual cages, weighed and fasted for 4h with free access to water. Blood glucose levels were measured from tail prick (Accu-check Performa glucometer; Roche, France) at basal (0min), 15, 30, 45, 60, 90 and 120min after oral administration of glucose (2 g.kg^-1^).

Locomotor activity and indirect calorimetry was performed in 8 mice of each group using a computer-controlled, open-circuit system (Oxymax Comprehensive Lab Animal Monitoring System; Columbus Instruments). Mice were individually tested during 24 h in clear chambers (20 × 10 × 12.5 cm) with a plastic elevated wire floor after being acclimated over 2 days. Room air was passed through the chamber at 0.6 l/min. The chamber exhaust was sampled for 45 s after an idle time of 90 s at 20-min intervals and was passed through O2 and CO2 sensors for estimation of oxygen consumption (VO2) and carbon dioxide production (VCO2). Outdoor air reference values were sampled every 8 measurements. Gas sensors were calibrated before experiments with gas standards containing known concentrations of O2, CO2, and N2 (Air Liquide—Certified accuracy ±1%). Respiratory exchange ratio (RER) was calculated as the ratio of carbon dioxide production (VCO2) to oxygen consumption (VO2). At the end of the calorimetric measurements, lean body mass (LBM) and fat mass (FM) were measured by quantitative NMR (EchoMRI 500T) to adjust energy expenditure (calculated as heat production = (3.815 + 1.232 × RER) × VO2)) to metabolic body size (LBM + 0.2 × FM) according to the procedure reported by Even and Nadkarni (Even & Nadkarni, 2012).

### Odorants preparation

Three odorants were chosen for their different chemical structures: an aromatic ketone, acetophenone, a terpene, (+) carvone and an aliphatic ester, amyl acetate. They were diluted into water while solubilized by DMSO. 10µL of the diluted odorant solutions were deposited on filter papers contained in Pasteur pipettes.

### Food finding test

Olfactory acuity was evaluated by means of a buried food test (Yang & Crawley, 2009) which relies on the natural tendency of mice to use olfactory cues while foraging. Pieces of Leerdamer cheese (squares of 5mm) were used as the food cue. Animals were habituated individually to the testing cage (conventional mouse cages (33x19x13 cm) filled with 6 cm of bedding) for 1 hour the day before testing. During this phase and just after being placed back in their grouped cage, they were given a piece of cheese on the bedding. We controlled that each mouse ate the cheese during this habituation phase. On the next day, animals were fasted for 5 hours and placed in individual cages. In the first session a piece of cheese was placed on the surface bedding pseudo randomly just before the introduction of the mouse in the center of the cage. The time to reach and start eating the cheese was measured with a manual timer and used to assess mice motivation to reach a food item. Mice were prevented from eating the entire cheese to avoid satiation. The two remaining trials were performed the same way, except that the cheese was buried beneath 3 cm of bedding. Test ended once either the animal found the cheese or did not achieve to find it within 10 minutes. The shortest time to retrieve and eat the cheese between the two trials was used to assess olfactory performance. Two days later, mice were tested for their motivation to dig in the bedding. They were fasted for 5h and placed in the same individual cages used during the first trial day. Their behaviour was monitored to measure the time spent digging during a 10-min trial in absence of any food cue.

### Effect of food-related odours on feeding behaviors

Mice were assigned to either a standard chow diet or the HFHS diet for 8 weeks before conducting experiments. Body weight was recorded weekly to monitor progression. Following the dietary intervention, mice were housed in automated monitoring cages (HM-2 system, MBRose), which enabled continuous, high-resolution tracking of individual food and water intake. Via RFID identification, this system provides precise quantification of meal timing, duration, and intake per animal. A 1-week acclimatization phase preceded any behavioural testing to ensure baseline feeding patterns stabilization. All mice underwent a food intake assessment in response to an odour exposure. Animals were subjected to a 24-hour baseline recording under their respective usual diet, followed by a 24-hour test recording during which food was soaked in an odorant solution. Two different odorant solutions have been used. Bacon odorant solution was prepared using bacon dithiazine molecule (W401755, Sigma-Aldrich) diluted at 10-2M (for 1mL: 10µL bacon dithiazine, 90µL dimethyl sulfoxide (DMSO), 900µL of mQH2O). Peanut butter odorant solution was prepared using peanut butter Menguy’s Peanut Butter Creamy) diluted in mineral oil (330779, Sigma-Aldrich) (for 5mL: 0,05g of peanut butter, 5mL of mineral oil). 20mL of odorant solution have been used to odorize one feeder of either chow or HFD pellets of 100g. Odorized food was introduced 1 hour preceding the onset of the active phase. All recordings started at the same time of the day to ensure similar metabolic conditions of the animals. All food intake data were continuously recorded via the HM-2 LAB system, exported in Excel format, and analysed using MATLAB to examine odour-driven modulation of feeding behaviour across dietary conditions.

### Electro-olfactogram recordings (EOG)

EOG were performed as previously described (Merle et al., 2019). Briefly, olfactory epithelium was exposed after decapitating mice and cutting their head on a sagittal plane. Recordings were made with glass micropipettes of 6-8 µm diameter filled with a mucosal saline solution (MS; 45 mM KCl, 20 mM KC_2_H_3_O_2_, 55 mM NaCH_3_SO_4_, 1 mM MgSO_4_, 5 mM CaCl_2_, 10 mM HEPES, 11 mM glucose, 50 mM mannitol, pH 7.4, 350 mOsm adjusted with mannitol). Reference electrode (Ag/AgCl) was placed in the olfactory bulb. Signal was amplified by an Axoclamp 200B amplifier (Axon Instruments, Molecular Devices, USA) and monitored with Axoscope (Axon Instruments, Molecular Devices, USA). Odorant air puff stimulation was delivered using a pressure controller for 200ms at 200mL/min inside a constant humidified airflow (1000mL/min) blowing on the olfactory mucosa through a 7mm-diameter tube. Analysis were performed using customed Matlab routines. Signal amplitude measured for odorant free stimulations was subtracted from odorant elicited signals. Depolarization amplitude, depolarization speed, fast (from 90 to 50%) and slow repolarization (from 50 to 10%) speeds of EOG responses were measured.

### NSO cilia reconstruction

Septal olfactory epithelia were extracted from the nasal cavity in a saline solution. They were mounted in a perfused chamber and visualized using an Olympus BX51WI microscope (40X objective + extra 2X magnification) coupled to a camera. Images were analyzed with FIJI and OSNs’ cilia were reconstructed using the FIJI plugin Simple Neurite Tracer. Fluorescent OSNs (SR1 expressing cells) were counted on the septal organ area.

### Glomeruli surface measures

Mice were anaesthetized with a mix of ketamine/xylazine (150 mg/kg and 10 mg/kg bodyweight, respectively) followed by a transcardiac perfusion with 4% paraformaldehyde solution. Brains were dissected and included in agar blocks to be cut in a vibratome into 50µm-thick transverse slices. GFP signal was acquired using a fluorescent microscope at 20x magnification (Axio Imager 2, Zeiss and the AxioVision software).

### Metformin administration

Metformin an insulin-sensitizing agent was chronically administered to HFHS mice in a specific experiment at the active dose of 300 mg/kg/day (Sigma-Aldrich) in drinking water during 4 weeks starting after 8 weeks of HFHS diet exposure. Water consumption was measured biweekly during the whole time of exposure to ensure the correct quantity of drug ingestion.

### Statistics

Data are expressed as mean ± standard error of the mean (SEM). Two-way ANOVA with repeated measures were used to determine statistical differences between groups in time series and in dose response curve designs. Post hoc multiple comparisons were performed using Fisher LSD tests. Two factor analyses were performed using either Mann-Whitney tests in case of absence of normality or Student’s t tests. A probability value of maximum p<0.05 was used as an indication of significant differences (*: p<0.05; **: p<0.01; ***: p<0.001; ****: p<0.0001).

## Results

### HFHS feeding impairs olfactory performance

Olfactory performance of HFHS fed mice presenting metabolic dysregulation including increased glucose intolerance and increased fat mass despite a similar body weight (Figure S1) was evaluated using the buried food test. HFHS fed mice took longer to retrieve a piece of cheese hidden under the bedding (Figure 1A) despite presenting similar motivation to forage for food as evaluated in measuring the time to retrieve a piece of cheese visible on top of the bedding and the time they spent digging in their bedding in absence of hidden food (Figure 1B, C). To test whether the olfactory impairments can influence food intake behaviour, we monitored their food consumption in response to two food odorants: bacon odour and peanut butter odour. Both odorants induced an increase in food intake during the first hour of exposure in control groups under chow diet (Figure 2) but not at later time points after 6h or 12h (Figure S2). By opposition, in HFHS fed mice, bacon odour exposure did not induce any variation in food intake (Figure 2A). Surprisingly, during peanut butter odour exposure, HFHS fed mice reduced their food consumption (Figure 2B). These results highlight an impairment of olfactory-dependent increase in food intake after 8 weeks of HFHS consumption.

**Figure 1:**
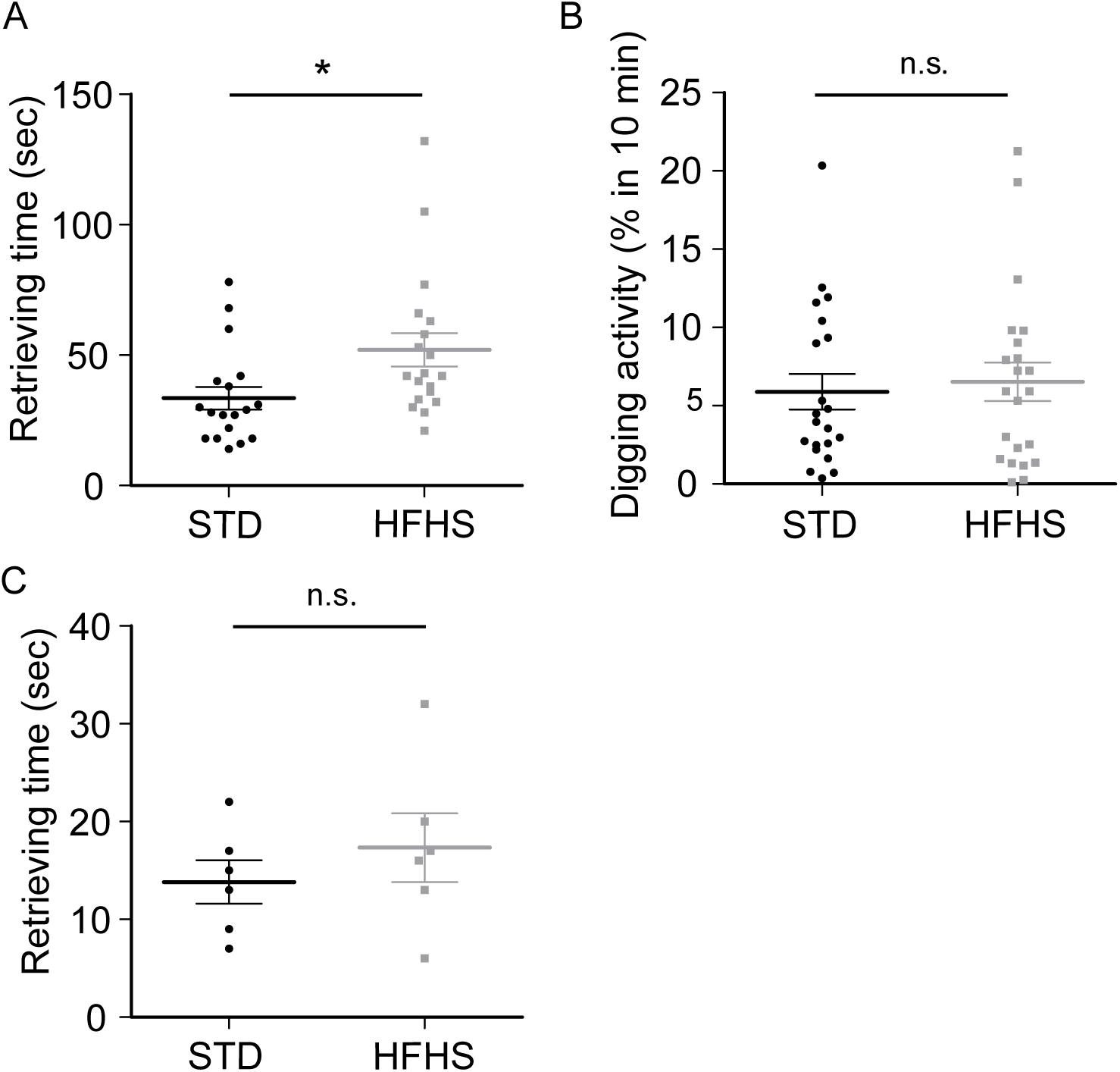
Buried food test in chow (STD) or HFHS fed male mice for 8 weeks. A. Retrieval time to find a cheese piece hidden under the cage bedding (STD: n=18; HFHS: n=19). Digging activity monitored during 10min in absence of any food in the cage, two days after the buried food test (B; STD: n=21; HFHS: n=22) and retrieval time for a piece of cheese visible on the bedding (C; STD: n=6; HFHS: n=6). HFHS animals showed a longer retrieval time than STD group. Control experiments (B, C) show no difference in their motivation to dig in their bedding Animals of the two groups showed similar times to retrieve cheese pieces when on the bedding. * p < 0.05; Student’s t test.

**Figure 2:**
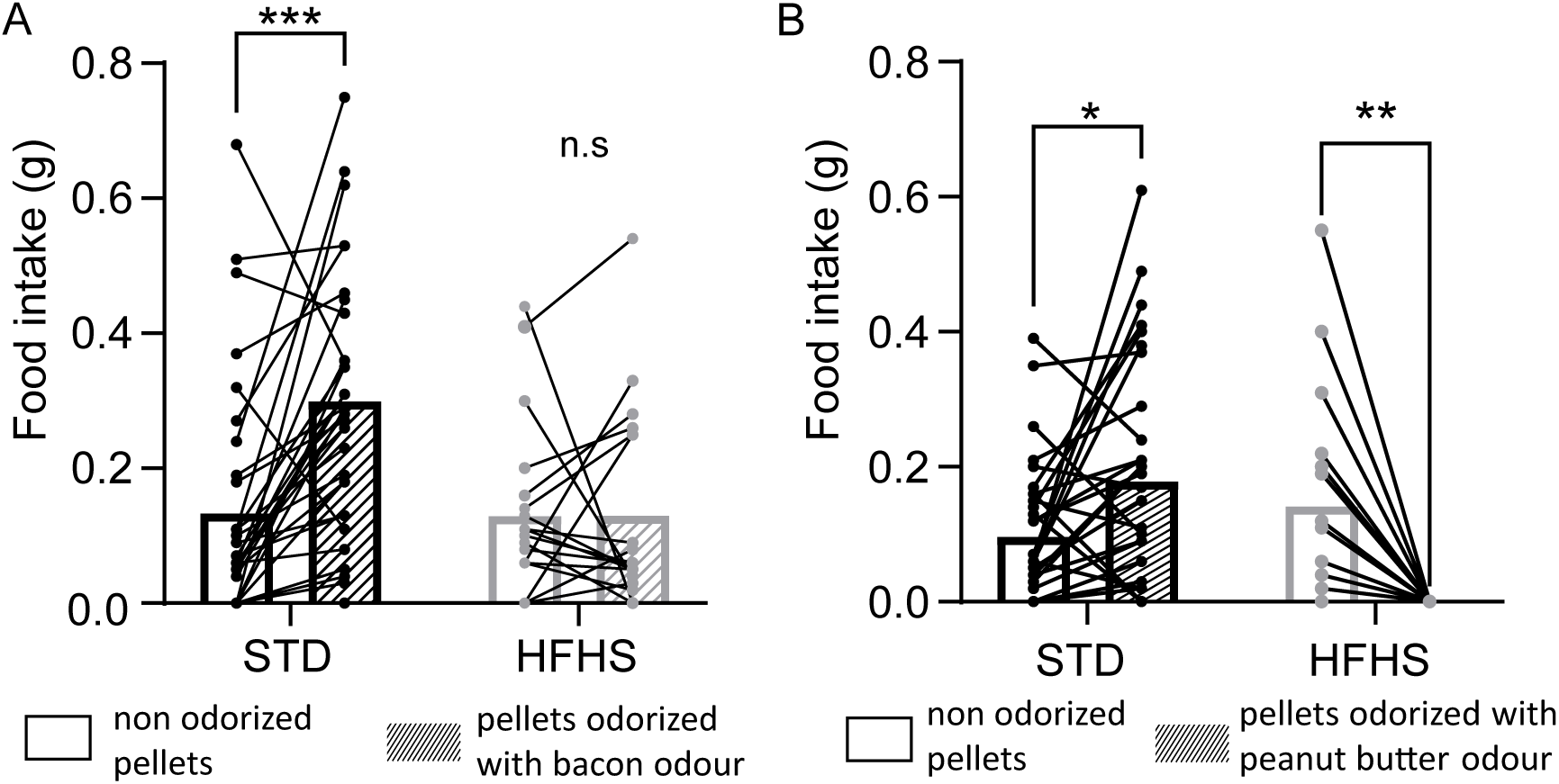
1h-food intake of odorized food in chow (STD) or HFHS fed male mice for 8 weeks. Consumption of food odorized with bacon odour (A; STD: n=32; HFHS: n=19) or peanut butter odour (B; STD: n=32; HFHS: n=19) during 1h after food access. Both odours increased the food intake of STD mice but not of HFHS mice. * p < 0.05; ** p < 0.01; *** *p* < 0.001; Student’s t tests.

### HFHS feeding alters olfactory mucosa

In order to investigate whether olfactory defects observed in HFHS mice are associated with alteration of the olfactory mucosa, we recorded the electrical response of a population of OSNs following odorant stimulations with EOG measurements. All tested odorants elicited sigmoidal dose-response relationships (Figure 3). EOG response amplitudes were lower in HFHS fed mice compared to control chow fed mice for acetophenone and amyl acetate and not statistically different for (+) carvone. These data show that olfactory mucosa functionality is impaired after 8 weeks of HFHS diet consumption.

**Figure 3:**
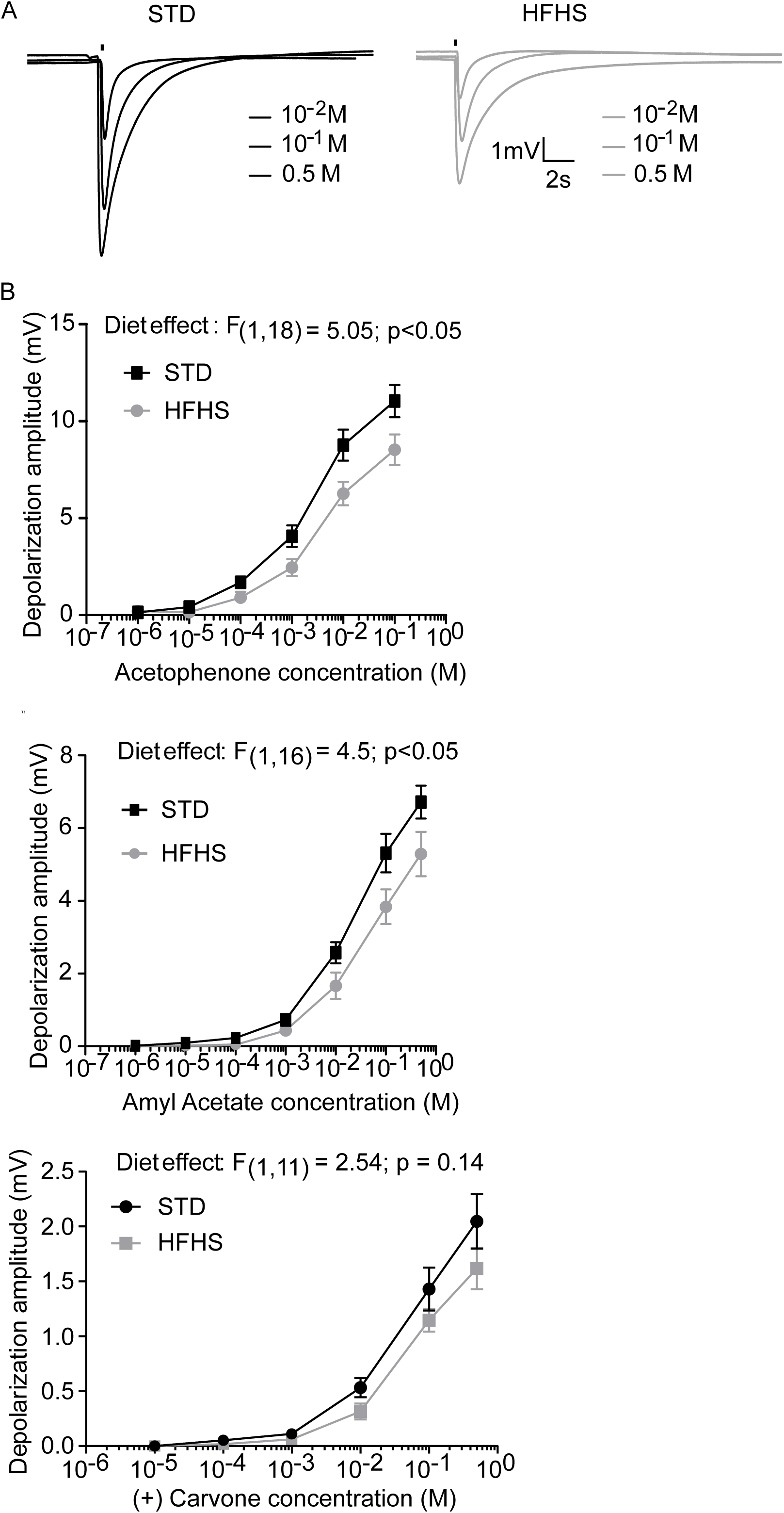
Electro-olfactogram recordings in chow (STD) or HFHS fed male mice for 8 weeks. A. Representative EOG traces recorded from the endoturbinate IIb in response to amyl acetate at 3 different concentrations applied during 200ms (black bar). B. Dose-response curves for acetophenone (STD: n=12; HFHS: n=8), amyl acetate (STD: n=9; HFHS: n=10) and (+) carvone (STD: n=6; HFHS: n=7). Recordings from HFHS animals showed lower amplitudes in response to varying concentrations of acetophenone and amyl acetate but not for (+)carvone. Two-Way ANOVA with repeated measures.

Reduced EOG response can indicate a smaller number of OSNs in the olfactory mucosa or a decreased detection of odours by individual OSNs. The use of a mouse line expressing GFP in specific OSNs expressing the SR1 olfactory receptor allowed to count these neurons in the olfactory epithelium and to reconstruct individual SR1-expressing OSNs (Figure 4). HFHS feeding did not impact the number of SR1-OSNs (Figure 4A, B). No difference between STD and HFHS fed mice was observed. The area of SR1-targeted glomeruli was quantified in coronal sections of the OB. HFHS feeding did not change the area of neither medial or lateral glomeruli (Figure 4E, F). However, the measurement of cilia length revealed that OSN cilia were shorter in HFHS fed mice compared to control chow fed mice (Figure 4C, D), supporting an impairment of OSN function as observed during the EOG.

**Figure 4:**
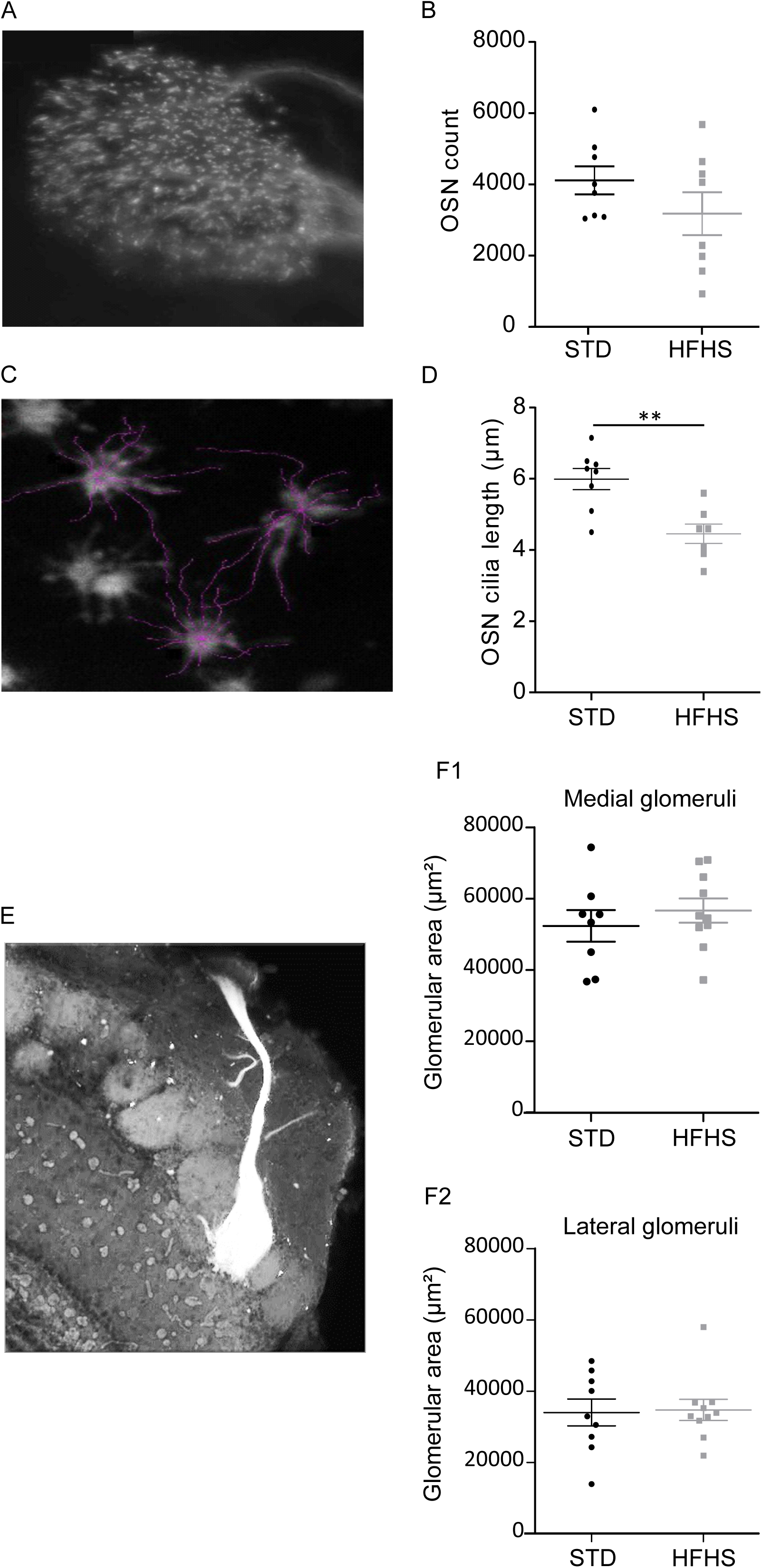
Impact of HFHS feeding on anatomical characteristics of SR1 expressing OSNs. Representative microphotograph (A) and quantification of SR1-GFP OSNs (B) in the septal organ in chow (STD, n=8) or HFHS (n=8) fed male mice for 8 weeks. Results show no difference in the number of OSNs. Representative microphotograph (C) and quantification (D) of olfactory cilia length from SR1-GFP OSNs with superimposed reconstruction in chow (STD, n=8) or HFHS (n=7) fed male mice for 8 weeks. Cilia were reconstructed using a semi-supervised method based on the GFP fluorescence. Cilia reconstructed from OSNs of HFHS mice were longer than STD group. Representative microphotograph of an olfactory bulb coronal section (E) and glomeruli area (F1, F2) in SR1-GFP mice fed chow (STD, n=8) or HFHS (n=10) for 8 weeks. Results show no difference in medial and lateral glomerulus sizes between the two groups. ** *p* < 0.01; Student’s t tests.

### HFHS withdrawal reverses olfactory deficits

We tested whether refeeding the HFHS group back with a standard chow diet would restore metabolic parameters and rescue the olfactory deficits. After 8 weeks of HFHS diet, a group of mice was fed a standard diet for 4 weeks (HFHS-STD) whereas a control group was maintained on standard diet for 4 additional weeks (STD-STD). 4 weeks after HFHS diet withdrawal to STD chow diet, all metabolic parameters were normalized back to controls (Figure S3). In the buried food test, no significant difference was observed in the time spent to retrieve the hidden piece of cheese between HFHS-STD and STD-STD fed mice (Figure 5A). Interestingly, HFHS-STD mice did not show any difference as compared to control STD-STD fed mice in the OSN cilia length (Figure 5B) or EOG response to acetophenone, amyl acetate or (+)carvone (Figure 5C). These data show that 4 weeks of standard chow diet were sufficient to restore metabolic parameters, olfactory mucosa functionality and olfactory deficits associated with HFHS feeding.

**Figure 5:**
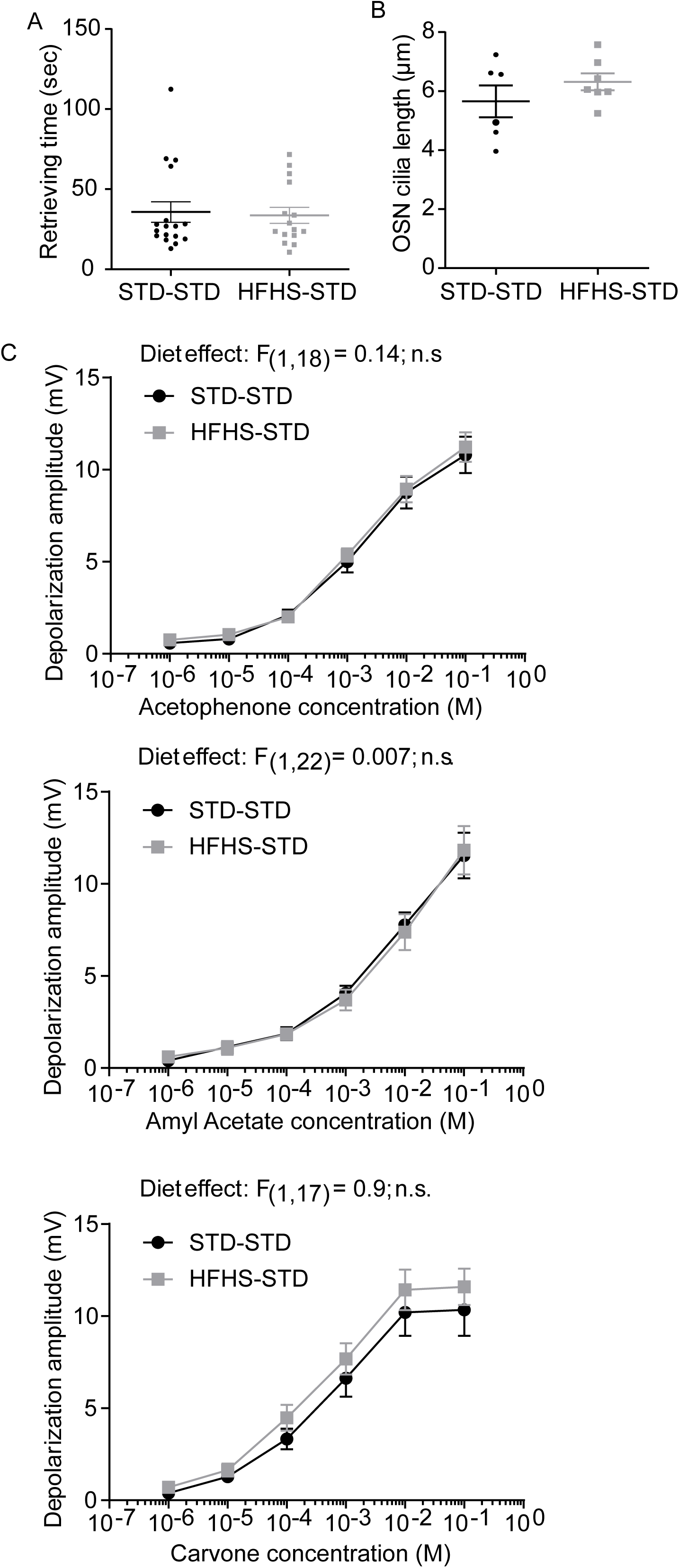
Diet reversal effect on metabolic and olfactory functions. SR1 male mice were fed a HFHS diet during 8 weeks and were later switched to a STD diet for 4 weeks (HFHS-STD). Results were compared to mice fed STD for 12 weeks (STD-STD). A. Time to retrieve a piece of cheese hidden under the bedding in the buried food test (STD-STD: n=22; HFHS-STD: n=19). Time was similar in both groups (Student’s t test: n.s.). B. OSN cilia length (STD-STD: n=6; HFHS-STD: n=7). No difference was observed between the length of OSN cilia measured in STD-STD fed mice and HFHS-STD fed mice. C. EOG dose-response curves for acetophenone (STD-STD: n=10; HFHS-STD: n=10), amyl acetate (STD-STD: n=12; HFHS-STD: n=12) and (+) carvone (STD-STD: n=9; HFHS-STD: n=10). Recordings from HFHS-STD and STD-STD mice showed similar amplitudes in response to the tested concentrations of acetophenone, amyl acetate and (+)carvone. Two-Way ANOVA with repeated measures: n.s.

Finally, we tested whether improving metabolic parameters with the antidiabetic metformin (MetF) while keeping animals under the hypercaloric HFHS diet would also improve olfactory capacities. Therefore, mice were fed for 8 weeks with a HFHS diet and treated for additional 4 weeks with MetF under the same HFHS diet. MetF treatment improved both glucose tolerance and basal blood glucose (Figure 6A, B). MetF treatment restored the EOG response to acetophenone (Figure 6C). However, MetF treatment did not restore EOG response to amyl acetate (Figure 6C) nor the olfactory sensitivity in the buried food test (Figure 6D). These data show that despite a potent restoration of metabolic parameters, MetF treatment only partially restored HFHS-induced olfactory impairments.

**Figure 6:**
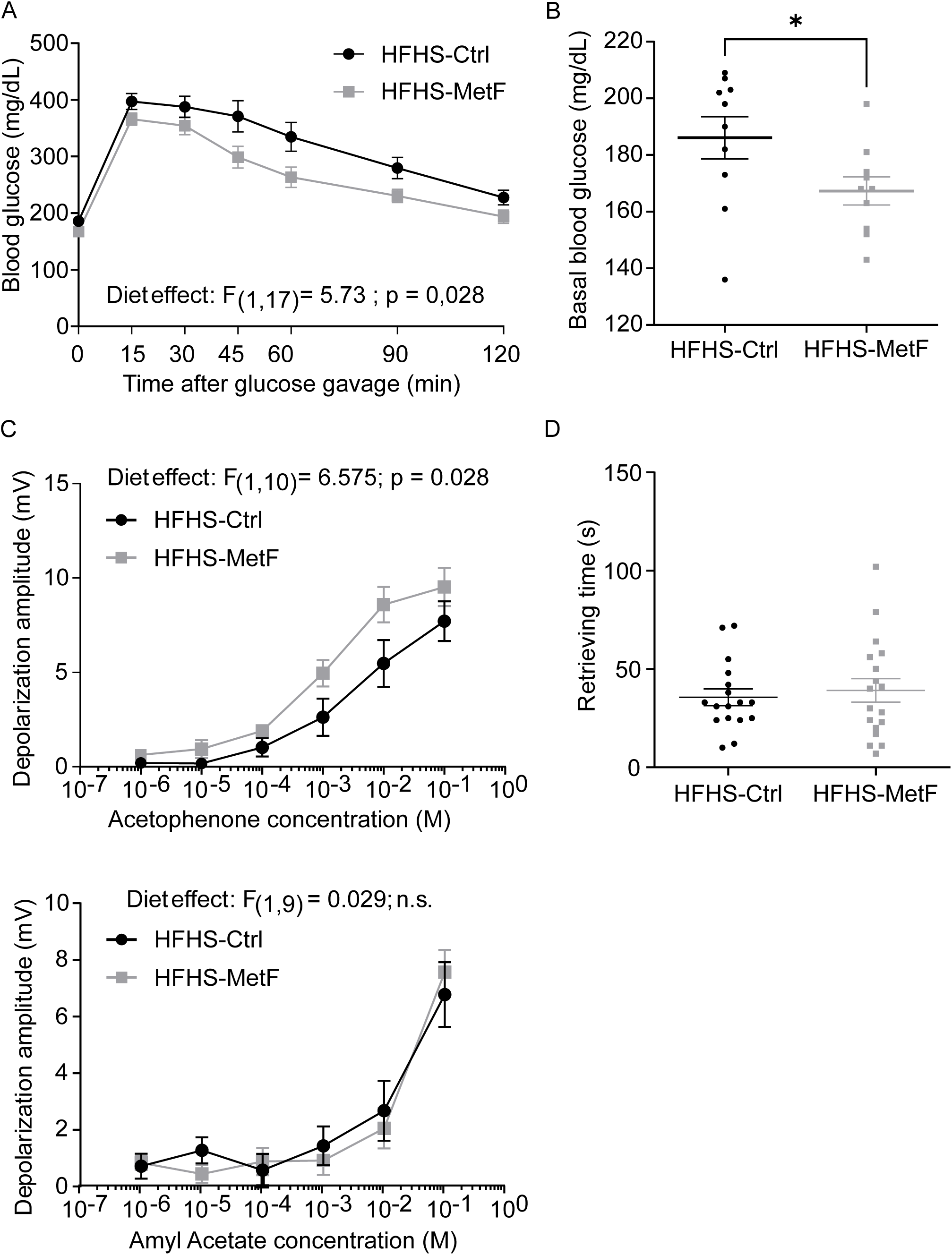
Effect of a chronic administration of metformin on metabolic parameters and olfactory capacities. Blood glucose level during an oral glucose tolerance test (2 g/kg; A; HFHS-Ctrl: n=10; HFHS-MetF: n=10) and basal blood glucose level (B; HFHS-Ctrl: n=10; HFHS-MetF: n=10) performed after 4 weeks of MetF administration in HFHS mice. HFHS-MetF animals showed improved glucose tolerance and basal blood glucose level. C. EOG dose-response curves for acetophenone (HFHS-Ctrl: n=6; HFHS-MetF: n=6), amyl acetate (HFHS-Ctrl: n=5; HFHS-MetF: n=6). MetF treatment improved the detection of Acetophenone in the HFHS-MetF group as revealed by the higher amplitudes of EOG responses compared to HFHS-Ctrl mice (Two-Way ANOVA repeated measures, p <0.05). No statistical difference was visible with amyl acetate stimulations (ANOVA repeated measures, n.s.). D. Buried food test. Time to retrieve a piece of cheese hidden under the bedding was similar in both groups (HFHS-Ctrl: n=17; HFHS-MetF: n=18; Student’s t test: n.s.).

## Discussion

Obesity has been associated with olfactory dysfunction in humans and in rodent models. Results presented here clearly showed that in absence of overweight the consumption of a high fat high sugar diet causes olfactory deficits. Olfactory mucosa, the site of odorant detection by OSNs is altered with both OSN morphology and functionality impaired. These alterations impacted the olfactory behaviour and modified the food intake response to attractive food cues. Reverting to standard diet restored OM functionality and olfactory capacities which could be explained by hormonal and structural changes as discussed below.

Numerous studies have reported the deleterious impact of diet induced obesity on olfactory capacities ((Challis et al., 2015)(Thiebaud et al., 2014); (Lietzau et al., 2020)). Overweight and obesity were postulated to be key in the development of olfactory deficits but subsequent studies including this one demonstrated the impact of hypercaloric feeding on the olfactory system independently from weight gain (Takase et al., 2016); (Chelette et al., 2022). Here electrophysiological measurements on the olfactory mucosa showed that reduced behavioural olfactory sensitivity observed in HFHS mice could originate from lower amplitude of odour responses in the olfactory epithelium. Even though the quantification of the number of OSNs did not reveal any difference between HFHS and STD mice, the shorter length of olfactory cilia could explain the lower mucosa sensitivity. The long distal segment of OSN cilia is the preferential site for the transduction cascade and signalling proteins such as ORs, G-proteins G_olf_, cyclic nucleotide gated channels and adenylate cyclase (Flannery et al., 2006; Takeuchi, 2024; Takeuchi & Kurahashi, 2018). Indeed, cilia increase the neuron’s ability to detect odor molecules by extending into the mucus and by increasing the surface area of the sensory neuron by as much as 40 times. Cilia length is a key component of OSN sensitivity to odorant molecules and a topographical graduation in cilia length has been exemplified in mouse olfactory mucosa that could match the odorant absorption in the nasal cavity (Challis et al., 2015). A reduction in the size of OSN cilia as observed in the HFHS diet condition of the present study is likely to decrease the efficiency of OSN activation upon odorant stimulation.

Impairments of the olfactory system could also be the consequence of changes in the metabolic status of HFHS fed animals (Palouzier-paulignan et al., 2012). Concordant with many other works HFHS diet led to numerous metabolic changes: increased fat proportion and epididymal fat deposits, impaired glucose tolerance without change in basal blood glucose. Despite a higher caloric intake in these mice, no overweight was observed after 8 weeks of HFHS diet exposure. Higher energy expenditure was measured in HFHS fed mice which has likely participated to the absence of weight gain as previously reported (Houtkooper et al., 2011). Fat accumulation has repeatedly been shown to increase the blood circulation of leptin (Frederich et al., 1995). Leptin receptors (Ob-R) are expressed in various cells of the olfactory epithelium including OSNs and their supporting cells (Baly et al., 2007; Getchell et al., 2006) and was indeed shown to be a potent modulator of the olfactory system (Palouzier-paulignan et al., 2012). Local administration of leptin reduced EOG responses in rat olfactory mucosa (Savigner et al., 2009). Other studies have documented the modulation of olfactory behaviour by leptin in rats. Leptin significantly decreased the time spent sniffing a food odour in fasted rats (Prud’homme et al., 2009) and reduced the sensitivity to an aversive odour (Julliard et al., 2007). The weaker EOG responses measured for acetophenone and amyl acetate in HFHS mice could derive from higher circulating leptin levels. This peripheral effect of leptin on the olfactory mucosa could explain the olfactory deficits revealed by the buried food test.

Insulin’s action on the olfactory mucosa shares a number of characteristics with leptin’s action. Insulin receptors are present in OSNs and other cells of the olfactory mucosa (Lacroix et al., 2008; Marks et al., 2009). This hormone showed the potential to modulate both the olfactory behaviour and the electrophysiological responses of OSNs (Lacroix et al., 2008; Marks et al., 2009; Savigner et al., 2009). Local administration of insulin reduced EOG responses in rat olfactory mucosa. Thus, decreased in insulin sensitivity suggested by the impaired glucose clearance during the oGTT could also take part in HFHS diet-induced decreased olfactory performance. Nevertheless, while the administration of the insulin-sensitizing agent metformin improved their metabolic parameters including glucose tolerance, this treatment only partially restored the EOG response and did not change the olfactory capacities measured in a buried food test. This would suggest that changes in insulin sensitivity could only partially explain HFHS-induced olfactory performance deficits. Finally, other metabolic peptides and hormones known to modulate the olfactory system among which orexigenic molecules such as peptide NPY (Negroni et al., 2012) or adiponectin (Loch et al., 2013) and modulated by HFHS feeding could take part in the observed alterations of the olfactory system.

Interestingly, HFHS impact on OSNs was reversible after refeeding animals standard chow diet. Both the morphology and the functionality of OSNs were restored by changing from a HFHS diet back to a STD diet. This nutritional modification resulted in a reduction of body fat mass and an improvement of glucose tolerance. This normalization of the metabolic parameters could be the cause of the recovery of the olfactory function after 4 weeks of standard chow consumption. Thiebaud et al (2014) in a comparable study reversed the diet back to chow after a 5 month-administration of a 60% fat diet and did not observed any evidence of reversibility of the olfactory function in spite of a weight and basal glycaemia normalization. Bariatric surgery in humans (particularly the sleeve gastrectomy) was shown to be effective in reversing olfactory decline associated with obesity (Peng et al., 2019). This surgical procedure leads to a drastic loss of weight accompanied by a normalization of the metabolic parameters.

A question which needs to be answered is to understand how nutritional changes impact OSN morphology including cilia length. Olfactory cilia share common characteristics with primary cilia i.e. microtubular architecture, enrichment of various receptors and other ciliary exclusive proteins (McClintock et al., 2020; Xie et al., 2022). Studies on ciliopathies have shown that maintenance of cilia architecture including the one of olfactory cilia requires a precise phospholipid composition (Xie et al., 2022). Primary cilia length and organisation was shown to be disrupted under diet induced obesity with shorter primary cilia observed in hypothalamic neurons (Han et al., 2014; Ritter et al., 2018). Fatty acid profile in olfactory cilia membranes was proven to vary according to the lipid composition of the diet (Soubeyre et al., 2023b). The HFHS diet used in the present study contained a high quantity of saturated fatty acids and presented a n-6/n-3 ratio of PUFAs equal to 10. As previous work has shown, olfactory mucosa contains a high proportion of n-3 PUFAs (Russell et al., 1989) and this composition can be altered by the diet (Khoury et al., 2020; Le Bon et al., 2018; Soubeyre et al., 2023a). Administration of the HFHS diet for 8 weeks could have altered the lipid composition of membranes of OSN cilia and therefore impacted cilia length and the overall functionality of OSNs.

Given the intricate relations between olfaction and nutrition, olfactory loss is expected to affect the feeding behaviour. Studies in humans showed that a gradual olfactory loss led to changes in dietary habits that ultimately produced either weight loss or weight gain (Aschenbrenner et al., 2008). Loss of smell can indeed decrease the hedonic sensations provided by food flavours and therefore reduce the pleasure to eat and the food intake. Olfactory loss could then act as a protective mechanism against further weight gain (Rasmussen et al., 2018). A study in mice showed that even short-term olfactory deficits could decrease food consumption and contribute to weight loss (Riera et al., 2017). However, another study revealed that anosmia did not impact the feeding behaviour and did not prevent the HFD induced weight gain (Boone et al., 2021). In the present study, mice under HFHS diet presented olfactory deficits and did not increase their food intake in response to the appetitive odours of bacon and peanut butter as STD mice did.

Data of the present study emphasizes the fact that overweight is not the cause of alterations of the olfactory system and further studies should be performed in human to verify this fact. Our results show structural changes in olfactory cilia that could be at the origin of the deficits of odour detection. Knowing the importance of olfaction in the regulation of feeding behaviour, the early loss of olfactory abilities due to Western diet consumption could influence food choices and intake.

## Aknowledgement

ANR StriaPOM (XF, DJ), GPR Brain NeuroFood (XF, DJ), University of Bordeaux Neurocampus Graduate School (LE), Bordeaux NeuroCampus (DJ), Regional Council of Burgundy France (FABER and PARI Agrale) (DJ, XG), European Funding for Regional Economical Development (FEDER) (XG), CNRS (ATIP and ATIP plus) (XG). Authors would like to thank Anne Lefranc and the CSGA animal facility and Gregory Artaxet and the CIRCE (Behavioural Engineering Centre) facility of Bordeaux Neurocampus for excellent animal care.

## Author contributions

Designed Research: X.F., X.G., D.J.; Performed experiments: L.E., E.A., V.C., J.P., T.C., R.C., V.S., D.J. Analyzed data: L.E., E.A., V.C., J.P., T.C., R.C., V.S., D.J; Writing – Original Draft: L.E., X.F., D.J.; Writing – Review and Editing: X.F., X.G., D.J.,; Supervision and Funding Acquisition: X.F., X.G., D.J.

