## Supplementary material for "Western diet reversibly alters the olfactory mucosa and impairs the response to appetitive food cues": Suppl. Figure

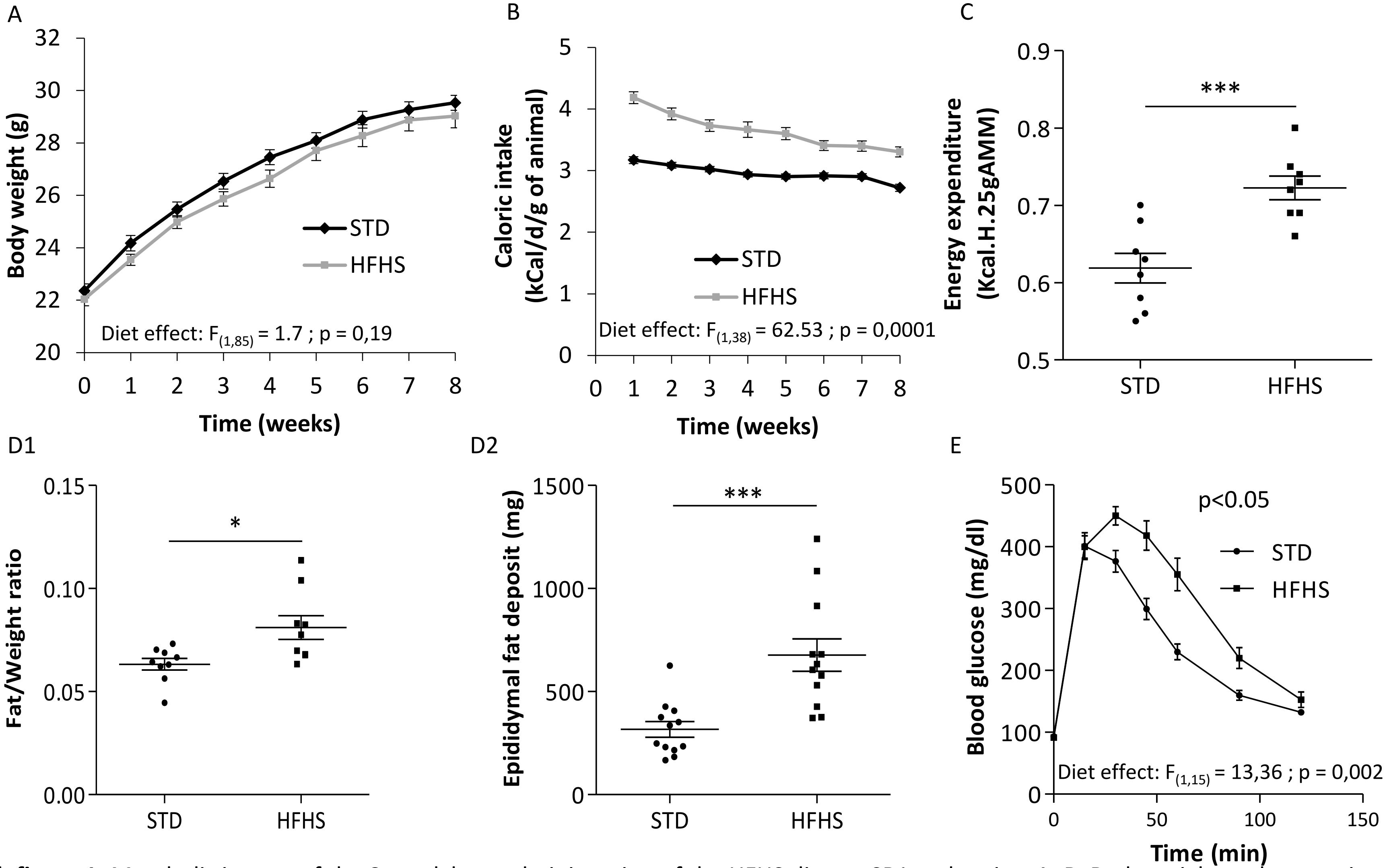


**Suppl. figure 1:** Metabolic impact of the 8-week long administration of the HFHS diet on SR1 male mice. A, B. Body weight and energy intake measured every week during the 8-week long administration period (mean ± sem; STD: n=44; HFHS n=43) Beside a higher energy intake (ANOVA repeated measures: $p<0.01$, HFHS animals showed no difference in weight gain compared to STD animals (ANOVA repeated measures: n.s.). C. Energy expenditure measured at the 8 week time point. HFHS animals used a larger amount of energy for a same activity rate. (mean ± sem; STD n=8; HFHS n=8; Student's t test: $p<0.001$) D. Fat accumulation measured by EchoMRI (D1) or as epididymal fat deposit (D2) after 8 weeks of diet. Both measures showed a higher fat mass in HFHS animals (mean ± sem; Student's t test: $p<0.05$ and $p<0.001$). E. Plasma glucose concentration over time in the oral glucose tolerance test performed after 8 weeks of diet administration. HFHS animals showed a longer time to recover their basal glycaemia after a glucose gavage (mean ± sem; STD n=10; HFHS n=10; Student's t test: $p<0.05$).

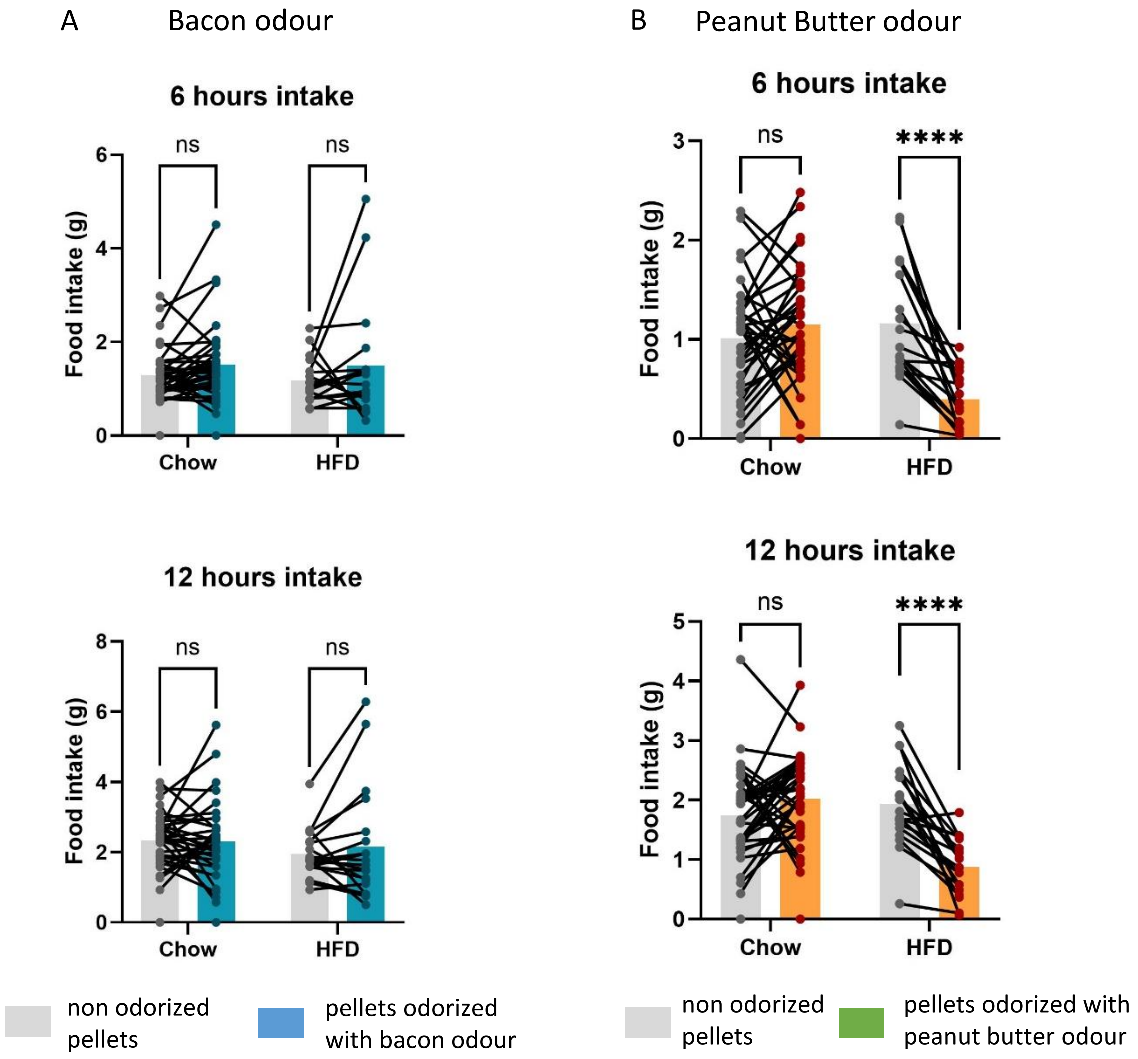


**Suppl. figure 2:** Consumption of food odorized with bacon odour or peanut butter odour in HFHS and STD fed mice. A. Consumption of food odorized with bacon odour during 6 and 12h after food access. Bacon odour did not change the food intake in both groups of mice at these time points. B. Consumption of food odorized with peanut butter odour during 6 and 12h after food access. HFHS mice decreased their food intake when pellets were odorized with peanut butter odour. Student's t tests: ***$p < 0.001$; **** $p < 0.0001$. Data are represented as mean ± sem.

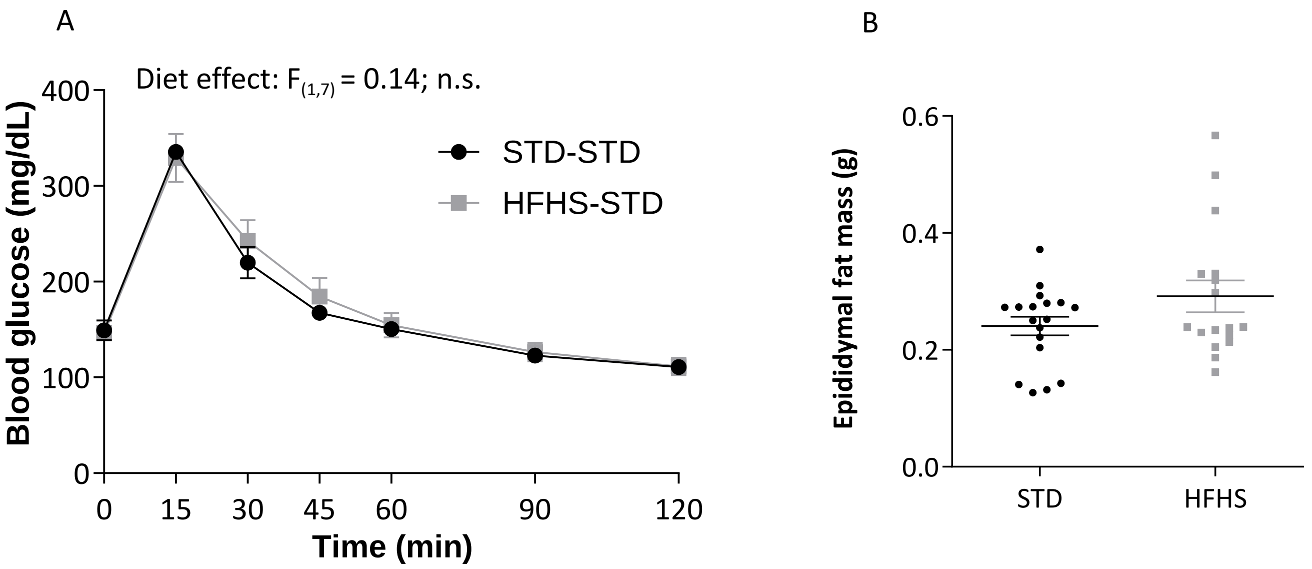


**Suppl. figure 3:** Diet reversal effect on metabolic parameters. SR1 male mice were fed a HFHS diet during 8 weeks and were later switched to a STD diet for 4 weeks (HFHS-STD). Results were compared to mice fed STD for 12 weeks (STD-STD). A. Plasma glucose concentration over time in the oral glucose tolerance test performed after 12 weeks of diet administration. HFHS-STD mice showed normal ability to recover their basal glycaemia after a glucose gavage (STD n=10; HFHS n=10; ANOVA repeated measures : n.s.). B. Epididymal fat mass measured in HFHS-STD and STD-STD mice. No difference was observed between the groups. (Mann Whitney test : n.s.)
